# Prior Novelty Invigorates Future Mesolimbic Target Detection

**DOI:** 10.1101/2024.12.16.628816

**Authors:** Blake L. Elliott, Kathleen J. O’Brien, Matthew Fain, Lauren M. Ellman, Vishnu P. Murty

## Abstract

The ability to adapt to a dynamic world relies on detecting, learning, and responding to environmental changes. The detection of novelty serves as a critical indicator of such changes, priming mechanisms to detect and respond to goal-relevant information. However, neural regions that support novelty detection (hippocampus) and goal-directed behavior (dopaminergic midbrain [VTA] and prefrontal cortex [PFC]) have yet to be described as a sequential process that unfolds over time. Using a forward-prediction functional magnetic resonance imaging (fMRI) model, we explored interactions between the hippocampus, VTA, and PFC in humans performing a novelty-imbued target-detection task. Hippocampal novelty activation predicted subsequent VTA target activation, enhancing readiness to detect goal-relevant information. Concurrently, goal-directed PFC activation modulated VTA target activation, refining focus on behaviorally significant cues. These circuits function both synergistically and independently, promoting subsequent hippocampal activity during target trials. This work provides new insights into how distributed circuits coordinate to optimize adaptive behavior.

**Significance Statement:** Surviving in dynamic environments requires coordinated neural mechanisms to detect, learn from, and respond to change. However, neural regions that support novelty detection and goal-oriented behavior have yet to be described as a sequential process that unfolds over time. Using a novel forward-prediction functional magnetic resonance imaging (fMRI) model, we show that hippocampal activation during novelty predicts ventral tegmental area (VTA) readiness to process goal-relevant information. Concurrently, goal-directed prefrontal cortex activity modulates VTA responses, sharpening focus on behaviorally significant cues. Furthermore, these synergistic and independent circuits enhance hippocampal sensitivity for future adaptive responses, offering novel insights into integration of brain mechanisms critical for learning, motivation, and executive function.

## Introduction

The ability to adapt to our dynamic world requires individuals to detect important features of new environments and respond appropriately. When you are in a novel environment, surrounded by unfamiliar sights and sounds, your attention naturally shifts toward meaningful cues (e.g., a sign for a taco shop if you are hungry). Importantly, novelty signals that the environment has changed, but it does not specify what the change is, when it is going to happen, or whether it is important. For an organism to be adaptive, novelty should prime and invigorate neural systems underlying the detection of behaviorally relevant information. In this sense, novelty may function as a gain modulator, enhancing the salience of motivationally relevant information while suppressing less pertinent information. While human and animal research has separately characterized circuits for novelty detection, target detection, and motivated learning, open questions remain about how these circuits interact to invigorate goal-directed behavior over prolonged timescales, when we may not know when the next important event is going to occur. Unlike associative learning wherein a cue would predict more information processing to an expected target, novelty should evoke a state of enhanced sensitivity to unpredictable targets. Here, we test a model by which neural regions that support novelty and goal-directed behavior (i.e., hippocampus, ventral tegmental area [VTA], and dorsolateral prefrontal cortex [dlPFC]), work in concert over time to invigorate mesolimbic circuits to respond to unpredictable goal-relevant information (i.e., target detection), even when it isn’t directly related to the preceding novel event.

Rather than producing a brief, time-locked neural response, novelty exposure has been shown to invigorate a broader neural state that primes the brain for flexible goal-directed behavior (Davis et al., 2004; Grace et al., 2007; Park et al., 2021; Lisman & Grace, 2005; Straube et al., 2003). Research in rodents has highlighted several circuits that are critical for translating novelty signals into goal-directed behavior. In their ‘hippocampal-VTA loop’ framework, Lisman and Grace (2005) propose a two-phase circuit: during the downward arc, novelty signals increase the gain of systems to promote goal-directed behavior, while during the upward arc, hippocampal plasticity is enhanced (Figure 1).

**Figure 1:**
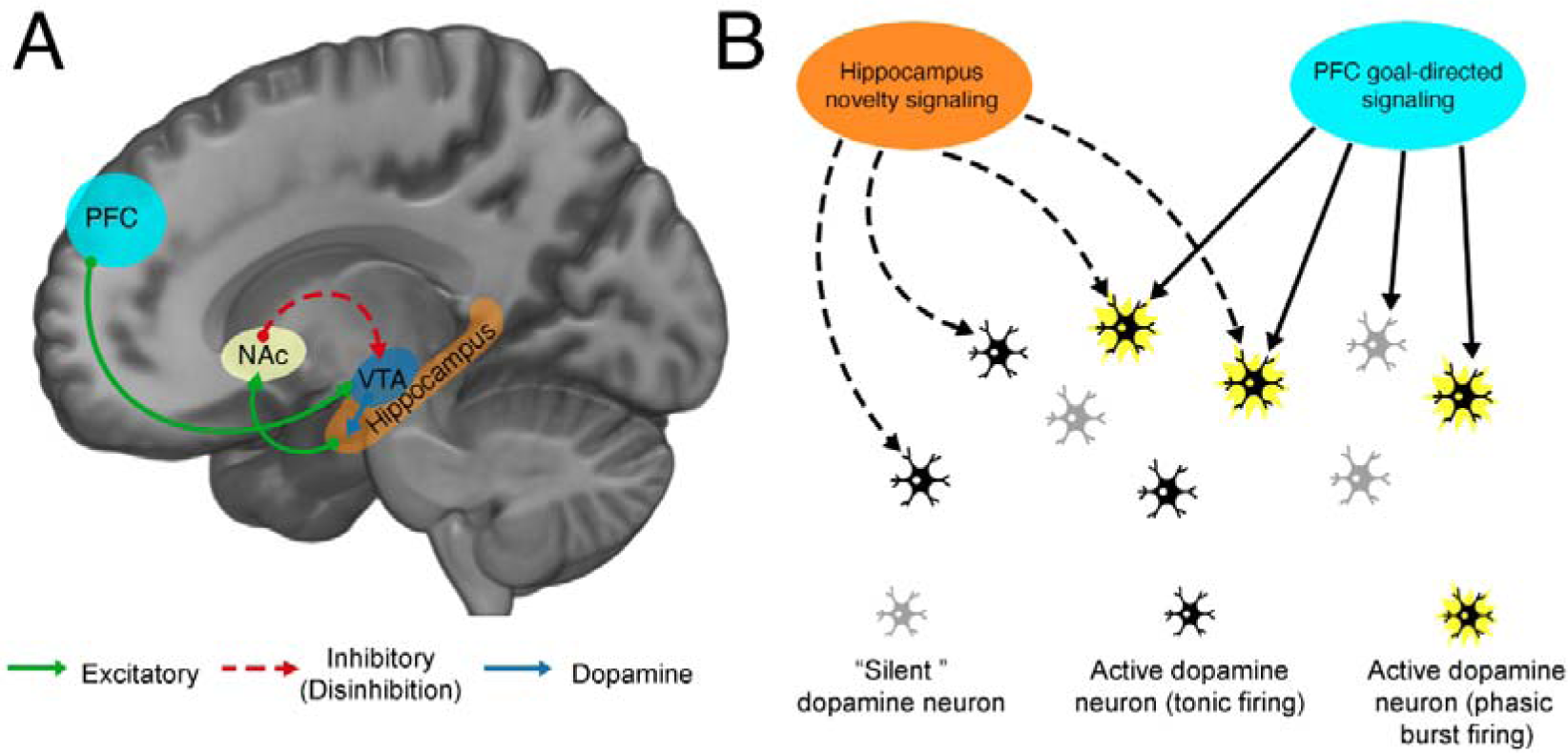
Afferent circuits modulating ventral tegmental area (VTA) activity. (A) The VTA is modulated by distinct afferent systems, the anterior hippocampus and prefrontal cortex (PFC). The anterior hippocampus modulates VTA activity via a polysynaptic circuit through the limbic striatum (i.e. nucleus accumbens [NAc]). The anterior hippocampus has an excitatory connection with the NAc, which forms a disinhibitory circuit with the VTA (via inhibition of the ventral pallidum, not pictured). The VTA has direct dopaminergic projections back to the hippocampus, influencing its function and stabilizing representations. The PFC has excitatory projections to the VTA. (B) VTA dopamine neuron activity states are modulated by the hippocampus and PFC. A substantial number of VTA dopamine neurons are not spontaneously firing (i.e. under tonic inhibition, “silent”). Hippocampal novelty signals release silent dopamine neurons from tonic inhibition, resulting in active spiking activity (i.e. tonic firing). In this way, hippocampal novelty signals control the baseline firing state of VTA dopamine neurons. In contrast, phasic burst firing, considered to be the behaviorally relevant activity of dopamine neurons, is modulated by the PFC. However, only neurons that are in a spontaneously active (tonic) state can elicit phasic burst firing. In this way, hippocampal novelty signaling and PFC goal-directed signaling synergistically modulate VTA dopamine neuron activity.

During the downward arc, the hippocampus detects environmental novelty (Bast, 2007; Kafkas & Montaldi, 2018; Kaplan et al., 2014; Knight, 1996), which in turn enhances tonic dopamine signalling in the VTA priming it to respond stronger to goal-relevant information when it occurs (Floresco et al., 2001, 2003; Legault & Wise, 2001). These tonic VTA signals are elevated for an extended period of time, invigorating the VTA to amplify incoming goal-relevant cues (Goto & Grace, 2007; Grace et al., 2007; Lodge & Grace, 2006a). In line with this model, prior work has shown that upon encountering a goal-relevant cue, the dlPFC, which is known to detect goal-relevant information (Friedman & Robbins, 2022a; Menon & D’Esposito, 2022; Miller & Cohen, 2001; Thompson-Schill et al., 2005), can stimulate phasic signaling in the VTA (Ballard et al., 2011; Carr & Sesack, 2000; Gariano & Groves, 1988; Murase et al., 1993). Phasic VTA signals, characterized by fast bursts of firing, mediate reward-related behaviors, including the refinement of goal-relevant responses (Adamantidis et al., 2011; Schultz, 2010, 2019; Stuber et al., 2008; Totah et al., 2013; Wanat et al., 2009).

During the upward arc, the synergistic engagement of the VTA by the hippocampus and dlPFC releases dopamine into the hippocampus, enhancing plasticity and subsequent memory formation (Huang & Kandel, 1995; Bethus et al., 2010; Davis et al., 2004; Frey & Morris, 1998; Gasbarri, Verney, et al., 1994; Redondo & Morris, 2011). Notably, in these paradigms, animals are first exposed to novelty, and then are exposed to unrelated, salient stimuli. Even though there is no predictive relationship between the two events, hippocampal novelty signals are still able to enhance mesolimbic processing of goal-relevant events (Li et al., 2003; Moncado & Viola, 2007; Ballerini et al., 2009; Wang et al., 2010). In this way, prior hippocampal novelty signals and concurrent dlPFC signals invigorate VTA responses and subsequent hippocampal dopamine. This last arm of the circuit dovetails with a larger literature showing VTA projections to the hippocampus enhances hippocampal sensitivity to and plasticity for new information, thus supporting long-term memory (Lisman et al., 2011; Lisman & Grace, 2005; Murty & Adcock, 2014; Shohamy & Adcock, 2010).

Much of this circuit has been explored piecemeal (or independently) in rodents via tracing (Carr & Sesack, 2000; Gasbarri, Packard, et al., 1994; Gasbarri, Verney, et al., 1994; Swanson, 1982), lesion studies (Gasbarri et al., 1996; Kelley & Mittleman, 1999; Lipska et al., 1992; Schmelzeis & Mittleman, 1996), and pharmacological manipulations (Floresco et al., 2001; Legault & Wise, 2001; Lodge & Grace, 2006a, 2008), which has limited the ability to understand the dynamics of this system in real-time. Human neuroimaging has also characterized independent parts of this circuit piecemeal during awake behavior, including novelty-evoked hippocampal signals predicting VTA tonic signalling (Murty et al., 2017), dlPFC interactions with phasic VTA signaling during goal-relevant behaviors (Ballard et al., 2011; Murty et al., 2017), and VTA interactions with the hippocampus during goal-relevant memory encoding (Miendlarzewska et al., 2016; Shohamy & Adcock, 2010). However, research has yet to investigate the downward and upward arc in tandem, precluding the ability to understand how and when the dynamic relationship between novelty-induced invigoration and goal-directed behavior emerges.

Traditionally, fMRI explores neural circuits using functional connectivity methods that look at coordinated engagement occurring synchronously. However, these traditional approaches are insufficient to capture how hippocampal-novelty signals promote states of enhanced target detection, as the novel events occur before target events. Following prior work (Murty et al., 2017; Cowan et al., 2021), we developed a new technique wherein we see how processing one event (i.e., novelty), precedes and predicts a subsequent goal-relevant event (i.e., target detection), wherein interactions across time reflect the asynchronous nature of novelty invigorating systems for goal-directed behavior (i.e., target detection).

The current study investigates hippocampus, VTA, and dlPFC dynamics in humans during a novelty-laden target detection task, in which target events occurred after viewing a series of scenes that vary in their relative composition of novel and familiar scenes, using fMRI. Although fMRI cannot directly measure dopaminergic signals, it affords the ability to examine multiple brain regions in awake humans during dynamic task performance. In line with the proposed downward arc of the hippocampal-VTA loop, we hypothesized that preceding hippocampus novelty activation will dynamically predict upcoming goal-directed VTA activation during target trials. Furthermore, we predicted that goal-directed behavior (i.e., target detection) would engage the PFC and be associated with increased VTA activation. In line with the upward arc of the loop, we hypothesized that simultaneous engagement of hippocampus-VTA-mediated novelty and PFC-VTA-mediated goal-directed circuits would predict hippocampus engagement to goal-relevant information (i.e., target detection).

## Materials and Methods

### Participants

The study was approved by Temple University’s Institutional Review Board. Participants were recruited for this experiment as healthy control subjects in a larger study examining psychosis risk. The final sample with usable task data and structural scans included 77 healthy, right-handed participants. Informed consent was obtained from each participant in a manner approved by Temple University’s Institutional Review Board.

### Procedure

The protocol and materials used in this experiment were based on previously published work (Murty et al., 2013, 2017). In brief, the task involved two phases: a familiarization phase and a novelty exposure phase (Figure 2). Approximately one hour before scanning, participants completed the familiarization phase, during which 120 outdoor scene images were presented sequentially (2 s each), followed by a prompt asking, “Have you seen this picture before?” Participants performed a self-paced continuous recognition task, pressing one of two buttons to indicate whether each image was “new” or “recognized.” Eighty images were repeated six times (“familiar”), while forty were presented once (“foils”). The familiarization task lasted approximately 20 minutes and was intended to establish a well-learned set of familiar scenes for subsequent testing. Roughly twenty minutes later, participants entered the MRI scanner for the novelty exposure phase. During this phase, participants viewed a randomized sequence of outdoor scene images, including 80 novel images (never seen before), 80 familiar images (from the prescan familiarization phase), and 40 presentations of a single target scene. Participants were instructed to press a button whenever the target scene appeared. Each trial consisted of a 2 s image presentation followed by a fixation cross that varied from 0.5 to 7 s (mean = 2.5 s), resulting in a total task duration of approximately 12 minutes 2 seconds. Importantly, participants were not informed that images from the familiarization phase would reappear, ensuring that novelty and familiarity were incidental rather than explicitly task relevant. All images were outdoor landscapes of equal visual complexity (100 ppi resolution), and image sets were counterbalanced across participants so that each scene appeared equally often as novel or familiar across subjects. Trial order and interstimulus intervals were optimized using OptSeq (Dale, 1999) to maximize estimation efficiency and minimize predictability. All trials were presented in a randomized order.

**Figure 2:**
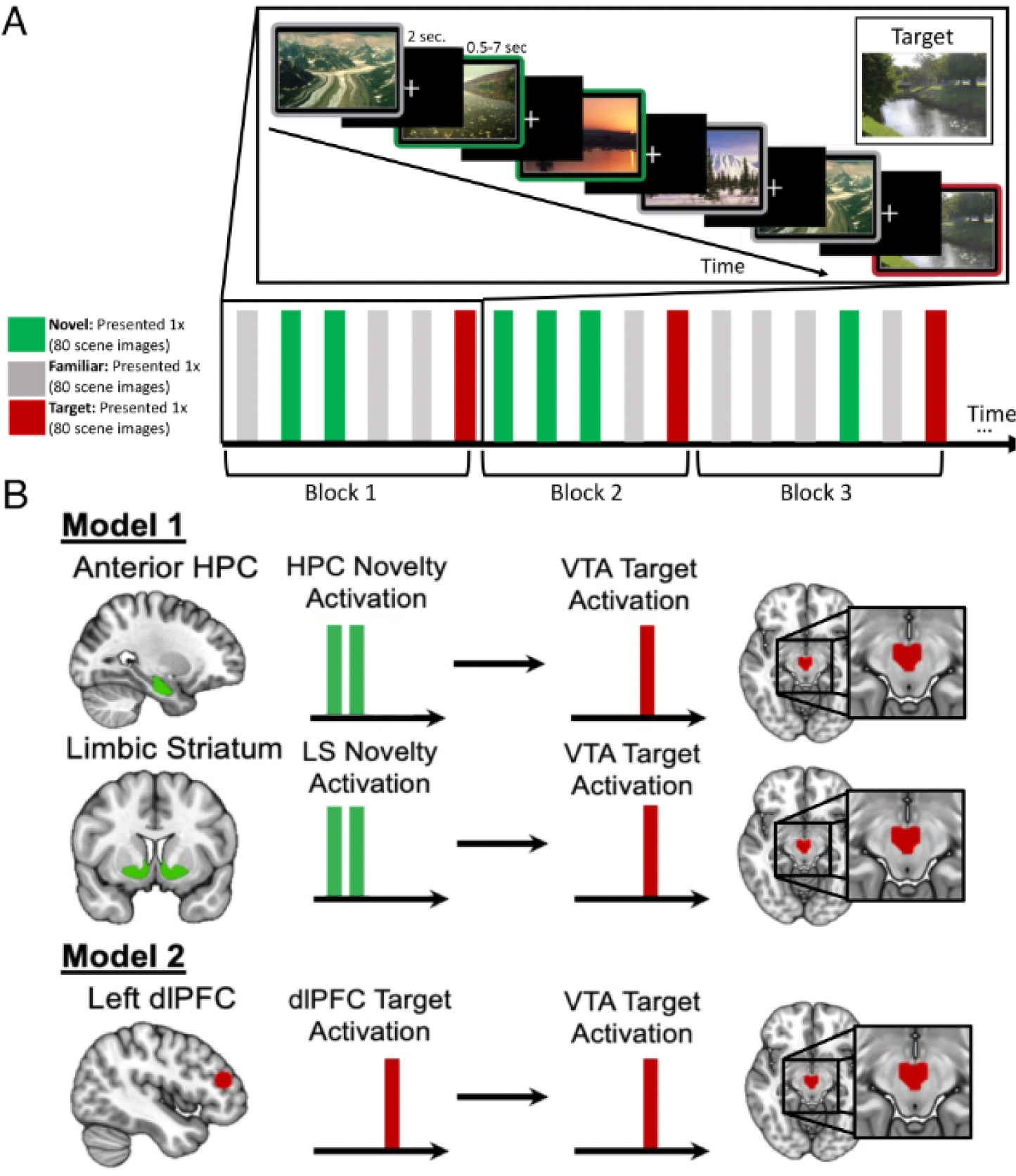
Experimental paradigm and fMRI model. (A) During the fMRI scan, participants performed a target detection task, pressing a button for a specific target image while incidentally viewing 80 familiar scenes (previously seen in the familiarization task) and 80 new scenes. (B) Forward-prediction fMRI models. Activation during novel and target trials was extracted on a segment-by-segment (block) basis. Linear mixed effect models were implemented to predict target VTA activation from preceding hippocampal (HPC) novelty and limbic striatum novelty activation (Model 1) or concurrent dlPFC target activation (Model 2).

### MRI Data Acquisition and Preprocessing

Scanning was performed on a 3T Siemens Magnetom Prisma scanner. Functional images were acquired using a multiband EPI sequence (TR = 1.73 s, TE = 25 ms, voxel size = 2.38 × 2.38 × 2.5 mm, multiband factor = 2). A high-resolution T1-weighted anatomical scan (MPRAGE; voxel size = 0.9 mm isotropic) was also collected to aid in functional image coregistration. Higher multiband acceleration factors (e.g., 4 or 8) have been shown to reduce task-related fMRI effect sizes in reward-related regions such as the nucleus accumbens (Srirangarajan et al., 2021). The chosen multiband factor of 2 provided a balance between increased temporal sampling and preservation of signal quality in reward-related regions.

Digital Imaging and Communications in Medicine (DICOM) images were converted to Neuroimaging Informatics Technology Initiative (NIFTI) format using dcm2niix (X. Li et al., 2016) and organized according to the Brain Imaging Data Structure (BIDS; (Gorgolewski et al., 2016)) using HeuDiConv (https://github.com/nipy/heudiconv). fMRI preprocessing followed the same workflow as a prior study from our laboratory (O’Shea et al., 2022). For consistency across manuscripts, we largely reproduce that description here. All preprocessing was performed using the established fMRIPrep pipeline (Esteban et al., 2019).

The T1-weighted (T1w) structural image was corrected for intensity non-uniformity (INU) using N4BiasFieldCorrection (ANTs v2.2.0; Tustison et al., 2010) and served as the anatomical reference throughout. The T1w-reference was skull-stripped with BrainExtraction.sh (ANTs), using OASIS30ANTs as the target template (Avants et al., 2008). Tissue segmentation into white matter (WM), gray matter (GM) and cerebrospinal fluid (CSF) was performed using FAST (FSL 5.0.9; Zhang et al., 2001). Spatial normalization to MNI152NLin2009cAsym standard space was performed via nonlinear registration using antsRegistration (ANTs v2.2.0), based on brain-extracted T1w images.

Per-participant fMRI preprocessing began with generation of a reference volume and its skull-stripped version using a custom methodology in fMRIPrep. Head-motion parameters (transformation matrices, six rotation/translation parameters) were estimated using MCFLIRT (FSL 5.0.9; Jenkinson et al., 2002) before any spatiotemporal filtering. Slice-timing correction was applied using 3dTshift (AFNI; Cox, 1996; Cox & Hyde, 1997). Using estimated susceptibility distortion, a corrected EPI reference was calculated to achieve more accurate co-registration with the T1w-reference.

The BOLD reference was co-registered to the T1w-reference using FLIRT (Jenkinson & Smith, 2001) with the boundary-based cost function (Greve & Fischl, 2009), configured with nine degrees of freedom to account for residual distortions. The BOLD time-series were resampled into native space by applying a single composite transformation correcting for motion and distortion. These were also resampled to MNI space, producing preprocessed BOLD runs in both native and standard space.

Confounding time series were calculated based on the preprocessed BOLD time series. Specifically, the first six aCompCor components (CSF, WM, and combined CSF+WM) were extracted and included as physiological noise regressors. Motion-related confounds were also modeled: framewise displacement (FD) and standardized DVARS, along with spike regressors identifying outlier volumes exceeding 1.5 mm FD or 2.0 DVARS. FD was computed as relative root mean square displacement between affines (Jenkinson et al., 2002). Following standard fMRIPrep practice, these measures were used for scrubbing via inclusion in the regression model (Power et al., 2012, 2014). In addition, global signals from CSF and WM were included. Head motion estimates, as well as their temporal derivatives (Satterthwaite et al., 2013), were also included in the confound file. All resampling operations were performed in a single interpolation step by composing all relevant transformations (head-motion, susceptibility distortion correction, and co-registration to anatomical and output spaces). Gridded volumetric resamplings were performed using antsApplyTransforms with Lanczos interpolation (Lanczos, 1964) to minimize smoothing effects of other kernels.

### Region of Interest (ROI) Definition

Anatomical hippocampal ROIs were defined using the Harvard Oxford Subcortical Probabilistic Atlas, thresholded at 50%, and was divided into anterior and posterior based on the last slice in which the uncus was visible (y = -21 on the MNI template, (Poppenk et al., 2013). The VTA ROI was defined based on a probabilistic atlas, thresholded at 50% (Murty et al., 2014). This atlas was created by averaging 50 hand-drawn ROIs utilizing anatomical landmarks in the midbrain, which allowed for the separation of the substantia nigra and VTA (Ballard et al., 2011; Naidich et al., 2009). The limbic (ventral) striatum (encompassing the nucleus accumbens) ROI was defined using a connectivity-based segmentation atlas with subdivisions for sensorimotor, executive, and limbic regions (Tziortzi et al., 2014). These striatal subdivisions also likely reflect separable functional pathways from midbrain dopaminergic regions (Elliott, D’Ardenne, Mukherjee, et al., 2022; Elliott, D’Ardenne, Murty, et al., 2022). Additionally, the homogeneity of dopamine release measured with positron emission tomography (PET) in response to the administration of amphetamine is significantly higher within these connectivity-based subdivisions of the striatum compared with within-anatomic subdivisions (i.e., putamen, caudate, and nucleus accumbens). Next, the ROIs (hippocampus, VTA, and limbic striatum) were resampled to fit the dimensions of each of the functional dataset using AFNI’s 3dResample (Cox, 1996).

### fMRI Data Analysis

Detailed procedures and analysis methods for the fMRI data were employed to ensure rigorous and reproducible results. fMRI data were analyzed using AFNI version 24.0.06.

#### Univariate Analysis During Novelty Exposure Task

To measure BOLD response during the novelty exposure phase of the task, we computed a GLM with regressors for each condition (novel, familiar, target) for each block (40 of each). A block was defined as the trials preceding a target (Figure 2). Individual events were convolved with a double-gamma hemodynamic response (HDR) function. Noise-related measures were also added as additional nuisance regressors. Noise-related measures were computed for average signal in CSF and white matter masks (generated using FSL’s FAST segmentation tool), time points of excessive head motion (identified using FSL’s motion outliers tool), as well as the six head motion parameters and their first derivatives. The resulting contrasts were registered to standard MNI space, from which we then extracted the b parameters from each condition, (e.g., novel, familiar, target greater than baseline) for each block, for each participant. We examined univariate responses across our ROIs of interest, in the hippocampus, VTA, and limbic striatum, and dlPFC regions. Due to the small size of the VTA and evidence that spatial smoothing can bias reward-related brain activity, no spatial smoothing was applied (Sacchet & Knutson, 2013).

### Statistical Analysis

#### Effect of Prior Novelty on Reaction Times

To investigate the behavioral relevance of exposure to novelty we conducted an analysis of reaction times to the target stimulus. Prior studies conducted in animals suggest that encountering novelty, even when completely orthogonal to the task demands, invigorates goal-directed behavior (via increased tonic VTA engagement). To test this, we first computed the temporal lag of each target from the last novel stimulus encountered. For each target trial, we assigned a lag value (ranging from −1 to −5) indicating how many trials had passed since the last novel stimulus, with lower values indicating greater temporal proximity to novelty. We then tested whether closer temporal proximity to a novel stimulus predicted faster reaction times during target trials using a linear mixed effects model. Specifically, we fit a model in which reaction time was predicted by lag from the last novel stimulus. All models were computed on a trial-wise level and included ‘participant ID’ as a random effect to account for within-subjects variation in the data. Model fit comparisons were described using AIC and BIC. Likelihood ratio tests were conducted using the anova() function to determine whether the full model provided a significantly better fit than the baseline model.

#### Afferent Circuits Modulating Target VTA Activation

We next examined how the afferent circuits associated with temporally distinct state-based engagement (i.e., hippocampal preceding novelty and dlPFC concurrent goal-directed) affect VTA activity during target detection. First, we examined if hippocampus-VTA connectivity increases when encountering a novel stimulus (a critical first step in mechanistically investigating hippocampus -VTA invigoration). Next, we examined whether preceding activation during novelty trials in the anterior hippocampus and limbic striatum predicted activation during the subsequent target trial in the VTA. Given the polysynaptic nature of the hippocampus-VTA circuit, the limbic striatum was included to examine if the activation was driven by the hippocampus instead of the intermediate striatal region. Next, we examined whether concurrent dlPFC target activation was associated with VTA target activation within a block. We implemented generalized linear mixed effects models for these analyses. These models were implemented using the lme4 package in R (lmer; R v 4.2.1), using a model comparison approach. In our first model we examined if the independent variable of ROI (anterior hippocampus novelty activation and limbic striatum novelty activation, within-subjects variables) improved the fit of a base model which included only participant ID (between-subjects variable) using an omnibus chi-squared test. In our second model we examined if the independent variable of ROI (dlPFC target activation, within-subjects variable) improved the fit of a base model which included only participant ID (between-subjects variable) using an omnibus chi-squared test. Model fit comparisons were described using AIC and BIC. All models were computed on a trial-wise level and included ‘subject’ as a random effect to account for within-subjects variation in the data. Likelihood ratio tests were conducted using the anova() function to determine whether the full model provided a significantly better fit than the baseline model.

To examine the specificity of the effect of preceding novelty and concurrent target trial activation on VTA target activation, we conducted permutation tests by shuffling trial labels, generating a null distribution of model coefficients across 1,000 permutations. This approach allowed us to test whether observed relationships between hippocampal and dlPFC activations with VTA target responses exceeded chance levels. Following each permutation, we recalculated model parameters and compared the resulting distributions to our observed data. These analyses ensured that the predictive effects were not due to random associations but instead reflected reliable temporal dependencies.

#### Time-Dependent Modulation of VTA Responses by Prior Novelty

To assess whether the influence of hippocampal novelty signals on target-related VTA activation varied as a function of temporal proximity to prior novelty, we computed a lag value for each target trial (ranging from −1 to −5), representing the number of trials since the last novel stimulus. We then included this lag variable as an interaction term in a linear mixed effects model predicting VTA activation from hippocampal novelty-related activation. This allowed us to test whether the relationship between hippocampal novelty responses and subsequent VTA activation was modulated by the temporal distance from prior novelty. All models were computed on a trial-wise level and included ‘participant ID’ as a random effect to account for within-subjects variation in the data. Model fit comparisons were described using AIC and BIC. Likelihood ratio tests were conducted using the anova() function to determine whether the full model provided a significantly better fit than the baseline model.

#### Independence of Hippocampal Novelty and Goal-directed PFC Circuits

To investigate whether hippocampus novelty activation and target dlPFC activation independently modulates VTA activity we employed a linear mixed-effects model using the ‘lmer’ function from the ‘lmerTest’ package. Our analysis involved a full model that included both predictors, with a random intercept for participant ID to account for individual variability. To ensure that the predictors did not exhibit multicollinearity, we calculated Variance Inflation Factors (VIFs). VIF values close to 1 indicated that multicollinearity was not a concern. To confirm that each predictor contributed uniquely to the model, we compared the full model with two reduced models, each excluding one predictor at a time. The first reduced model excluded hippocampal novelty signaling, and the second excluded target dlPFC signaling. Model fit comparisons were described using AIC and BIC. Likelihood ratio tests were conducted using the anova() function to determine whether the full model provided a significantly better fit compared to the reduced models.

#### Hippocampal Long-axis Specificity

Animal models predict that ventral (anterior in humans) hippocampal novelty signals modulate the VTA. Thus, we tested whether anterior versus posterior hippocampal novelty signals predicted subsequent VTA activation using linear mixed effects models and a model comparison approach. To test if posterior hippocampal signals predicted subsequent VTA target activation, in our first model we examined if the independent variable of ROI (posterior hippocampus novelty activation, within-subjects variable) improved the fit of a base model which included participant ID (between-subjects variable) using an omnibus chi-squared test. To test if the posterior hippocampus better predicted VTA target activation than the anterior hippocampus we next examined if the independent variable of ROI (posterior hippocampus novelty activation, within-subjects variable) improved the fit of a base model which included participant ID (between-subjects variable) and anterior hippocampus novelty activation (within-subjects variable) using an omnibus chi-squared test. Model fit comparisons were described using AIC and BIC. All models were computed on a trial-wise level and included ‘participant ID’ as a random effect to account for within-subjects variation in the data. Likelihood ratio tests were conducted using the ‘anova()‘ function to determine whether the full model provided a significantly better fit compared to the reduced models.

#### Specific versus Relative Novelty

We next examined whether preceding hippocampal novelty activation predicting target VTA activation is specific to activation during the novel images only (specific novelty) or is sustained throughout the preceding block (relative novelty). We tested these hypotheses using linear mixed effects models and a model comparison approach. In our models we examined if the independent variable of ROI (hippocampal response to familiar scenes only [relative novelty], within-subjects variable) improved the fit of a base model which included participant ID (between-subjects variable) and hippocampal response to novel scenes only (specific novelty, within-subjects variable) using an omnibus chi-squared test. Model fit comparisons were described using AIC and BIC. All models were computed on a trial-wise level and included ‘participant ID’ as a random effect to account for within-subjects variation in the data. Likelihood ratio tests were conducted using the ‘anova()’ function to determine whether the full model provided a significantly better fit than the baseline model.

#### Afferent Circuit Modulation of Hippocampus Sensitivity

To investigate how hippocampal and prefrontal modulation of the VTA (i.e., the downward arc) contribute to subsequent hippocampal sensitivity during target trials (i.e., the upward arc), we implemented a path analysis using the lavaan package (Rosseel, 2012) with our time-lagged fMRI model. Regression coefficients were estimated using maximum likelihood, and mediation effects were calculated with the product-of-coefficients approach. Standard errors (SEs) and 95% confidence intervals (CIs) were estimated via bootstrapping (1,000 samples), which is robust to non-normality, increases power, and minimizes imprecision due to small sample sizes (Kelley, 2005). Global model fit was assessed using the comparative fit index (CFI) and root mean square error of approximation (RMSEA), with benchmarks of CFI > 0.90 and RMSEA ≤ 0.07 indicating good fit (Kline, 2015; Steiger, 2007). Pathways were considered significant if the 95% CI did not include zero. All coefficients are presented as standardized (β) values to allow comparison of effect magnitudes within and across models. Standardized path coefficients were used as path effect sizes (Nieminen et al., 2013).

## Results

### Temporal Proximity to Novelty Predicts Faster Target Responses

To test whether temporal proximity to novelty trials influenced goal-directed behavior (target-detection reaction time), we examined whether the temporal lag from the last novel trial predicted reaction time during target trials (*M* = 0.71s, *SD* = 0.24s). Confirming our hypothesis, we found that adding lag from the last novel trial as a predictor significantly improved model fit for predicting reaction time (AIC and BIC without lag: −270.36, −253.73; AIC and BIC with lag: −272.67, −250.49; model comparison: χ²(1) = 4.30, *p* = 0.04, Figure 3). The fixed effect of lag was significant, such that greater temporal distance from novelty was associated with slower reaction times (β = 0.010, *SE* = 0.0048, *t*(1811) = 2.08, *p* = 0.04). These results suggest that proximity to prior novelty facilitates faster behavioral responses (Figure 3).

**Figure 3.**
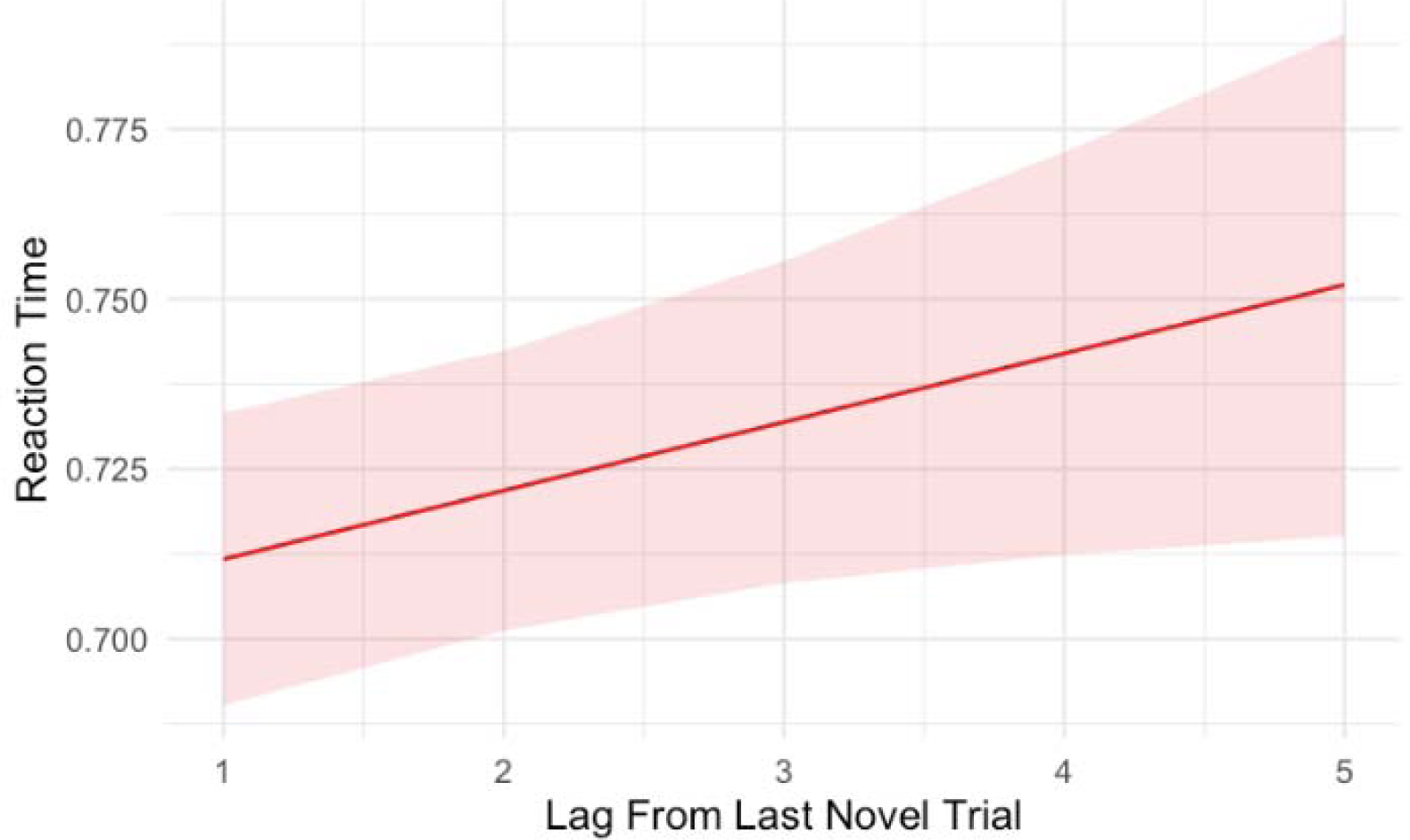
Reaction time (seconds) increases as a function of temporal lag from the last novel trial. Linear mixed effects modeling revealed a significant positive relationship between lag from the last novel stimulus and reaction time on target trials, such that greater temporal distance from novelty predicted slower responses. The shaded area represents the 95% confidence interval around the fitted regression line.

### Hippocampal-VTA Coupling During Novelty

Animal models predict that, in response to a novel stimulus, the anterior hippocampus (HPC) upregulates ventral tegmental area (VTA) activity through disinhibitory pathways. This model posits that such hippocampal-driven modulation of VTA should occur concurrently with the novel event. To test this prediction, we compared a baseline model including only subject-level random effects to a model that added hippocampal novelty activation as a fixed predictor of concurrent VTA activity. Adding hippocampal novelty activity significantly improved model fit (AIC and BIC without hippocampal novelty: –1675.8, –1659.0; AIC and BIC with hippocampal novelty: –1740.4, –1718.0; model comparison: χ²(1) = 66.56, *p* = 3.4×10^⁻¹^L), indicating that greater hippocampal activation during novelty trials robustly predicted greater concurrent VTA activation (Figure 4). These results provide strong evidence for time-locked hippocampal–VTA coactivation during novelty processing.

**Figure 4:**
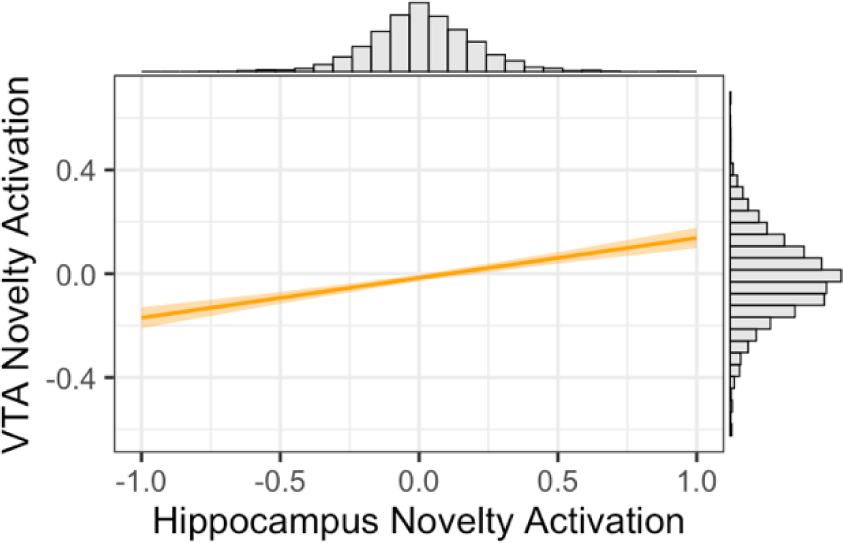
Time-locked hippocampus–VTA coupling during novelty. Predicted VTA novelty activation as a function of hippocampal novelty activation from a linear mixed-effects model (random intercepts for subjects). The solid line depicts the fixed-effect estimate; the shaded band is the 95% CI; marginal histograms show the distributions of each variable.

### Hippocampal Novelty Activation Predicts Target VTA Activation

To test whether hippocampal novelty signals affect subsequent goal-relevant phasic VTA responses, we investigated whether activation in the hippocampus during novel trials predicted subsequent event-evoked responses in VTA during target trials. Confirming our predictions, we found that adding hippocampal novelty activation as a predictor significantly improved model fit for predicting target VTA activation (AIC and BIC without hippocampal novelty activation: -1981.0, -1964.4; AIC and BIC with hippocampal novelty activation: -1987.1, -1964.9; model comparison: χ²[1] = 8.06, *p* < 0.01, Figure 5A). Given the polysynaptic nature of the hippocampal circuit through the limbic striatum, we next ran a model that included limbic striatum activation. The model comparison showed that including the limbic striatum did not significantly improve model fit compared to including only hippocampal novelty activation (AIC and BIC without limbic striatum activation: -1987.1, -1964.9; AIC and BIC with limbic striatum activation: -1986.8, -1959.1; model comparison: χ²[1] = 1.75, *p* = 0.186).

**Figure 5:**
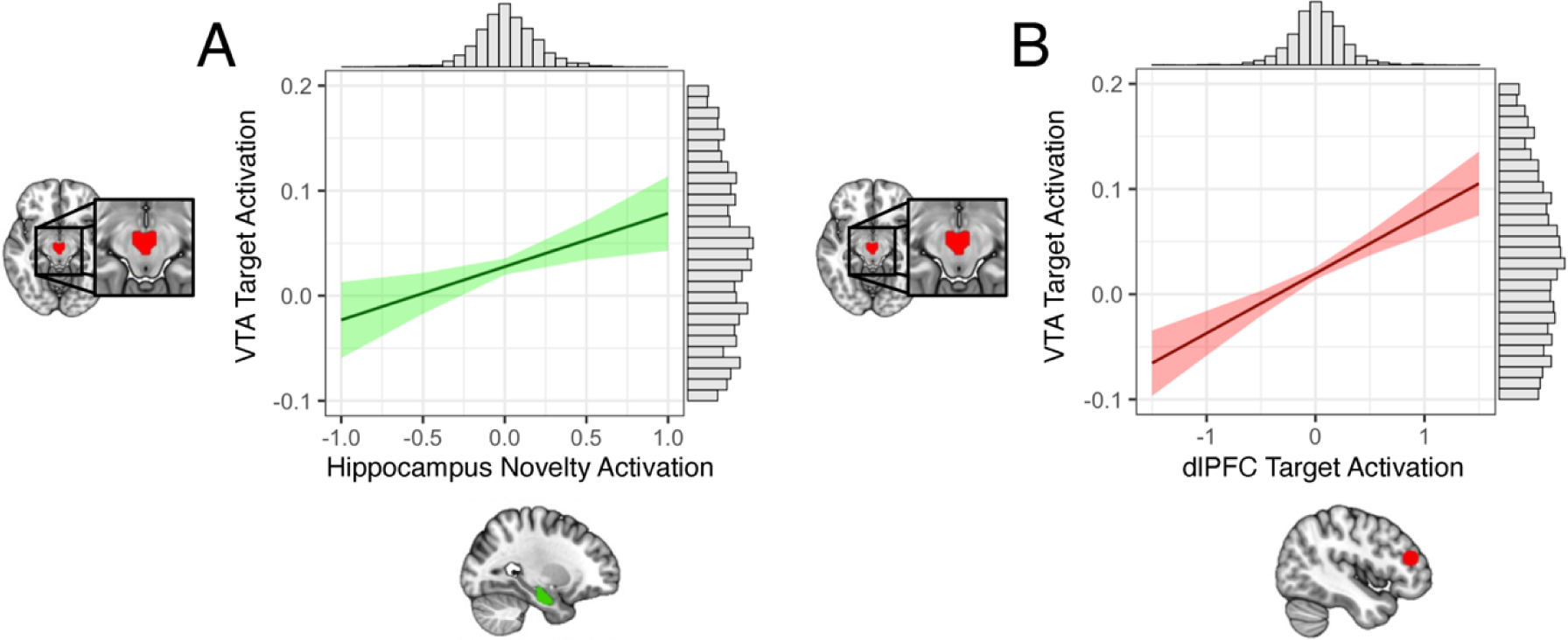
Estimated marginal effects from the linear mixed effects models. (A) Predicted values of VTA target activation as a function of hippocampus novelty activation, averaged over the random intercepts for subjects. The solid line represents the fixed effect estimate, and the shaded area indicates the 95% confidence interval. (B) Predicted values of VTA target activation as a function of dlPFC target activation, averaged over the random intercepts for subjects. The solid line represents the fixed effect estimate, and the shaded area indicates the 95% confidence interval.

We next tested whether hippocampal novelty responses predicting target VTA activity were temporally specific (i.e., the preceding novelty trials) by implementing a permutation test in which we shuffled the trial labels, generating a distribution of model coefficients (t-values) across 1,000 permutations. The confidence interval (CI) of the t-values for the model coefficients ranged from -1.61 to 2.44, indicating that the observed effects did not significantly deviate from those expected by chance at the 95% confidence level. This permutation test indicated that the effect of novelty on target VTA activation was specific to the preceding novelty trials from the same block (Figure 6A).

**Figure 6:**
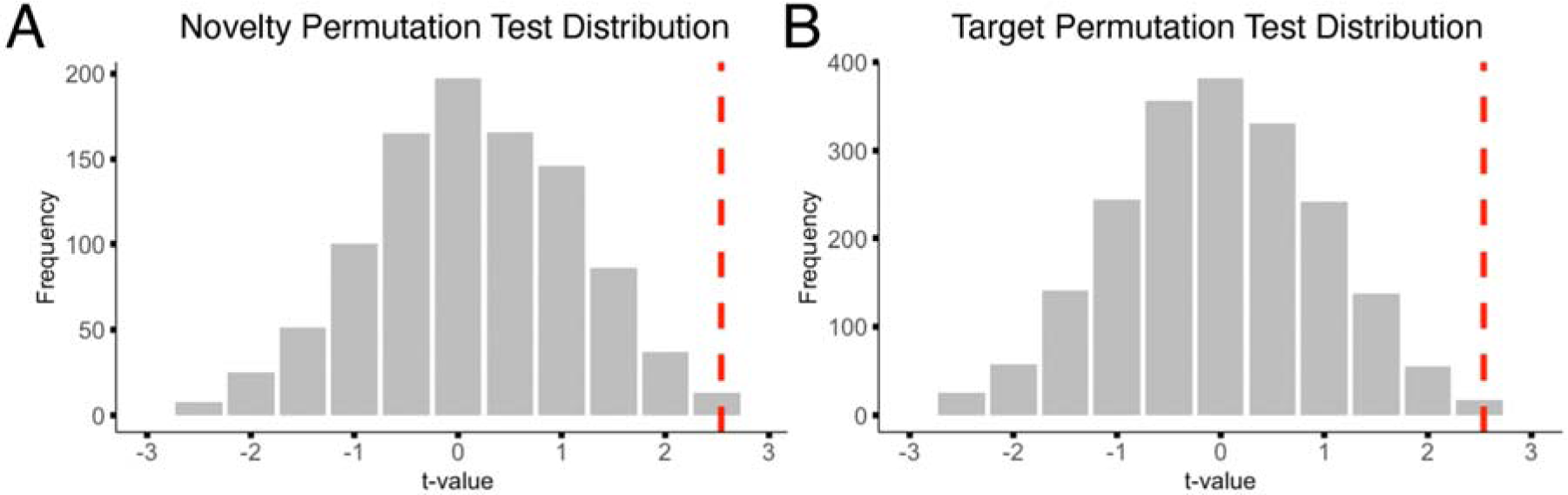
Permutation distributions. (A) Distribution of t-values from shuffling the trial labels of hippocampal novelty trials predicting target VTA trials. (B) Distribution of t-values from shuffling the trial labels of dlPFC target trials predicting target VTA trials. The dashed red line represents the observed t-value of the predictor from the model.

### dlPFC Target Activation Predicts VTA Target Activation

To test whether dlPFC target signals affect phasic VTA responses, we investigated whether activation in the dlPFC predicted phasic VTA responses during goal-directed behavior (target trials). Including dlPFC target activation in the model significantly improved fit over the baseline model (AIC and BIC without dlPFC target activation: -3240.4, -3222.3; AIC and BIC with dlPFC target activation: -3268.8, -3244.7; model comparison: χ²[1] = 30.44, *p* < 0.001, Figure 5B).

We next tested whether these effects were temporally specific by implementing a permutation test in which we shuffled the trial labels, generating a distribution of model coefficients (t-values) across 1,000 permutations. The confidence interval (CI) of the t-values for the model coefficients ranged from -1.79 to 2.19, indicating that the observed effects did not significantly deviate from those expected by chance at the 95% confidence level, indicating that the effect of target dlPFC activation on target VTA activation was specific to the target trials from the same block (Figure 6B). In accordance with the rodent literature, these results indicate that phasic dlPFC target activation predicts phasic VTA target activation.

### Independence of Hippocampal Novelty and Goal-directed PFC Circuits

We next investigated whether hippocampus novelty activation and target dlPFC activation independently influences goal-directed phasic VTA responses. We fit a full model that included both hippocampal novelty activation and target dlPFC activation as fixed effects, while accounting for individual variability with a random intercept for participant ID. Variance Inflation Factors (VIFs) were calculated to check for multicollinearity between predictors. Both VIF values were approximately 1 (Hippocampal Novelty: 1.002, dlPFC Target: 1.002), indicating that multicollinearity was not a concern. Examining the fixed effects in the full model, both predictors were statistically significant. Hippocampal novelty activation was a significant positive predictor of VTA activation, *t*(1864) = 2.61, *p* < 0.01, and target dlPFC activation showed a robust positive association, *t*(1865) = 5.51, *p* < 0.001.

Likelihood ratio tests revealed that removing hippocampal novelty activation significantly worsened the model fit, χ²(1) = 6.82, *p* < 0.01, suggesting that hippocampal novelty activation contributes uniquely to VTA engagement. Similarly, removing target dlPFC activation resulted in a significant reduction in model fit, χ²(1) = 30.12, *p* < 0.001, indicating that target dlPFC activation also independently predicts VTA engagement. These results support the hypothesis that hippocampal novelty and goal-directed PFC circuits independently contribute to VTA engagement.

### Hippocampal Long-Axis Specificity

Rodent research suggests that these effects would be specific to ventral regions of the hippocampus akin to the anterior hippocampus in humans (Fanselow & Dong, 2010; Poppenk et al., 2013; Strange et al., 2014). Therefore, we examined whether the observed effects were specific to the anterior hippocampus. We first tested whether including posterior hippocampal novelty activation would improve model fit compared to a baseline model. Interestingly, we found that including posterior hippocampal activation in the model significantly improved fit over the baseline model (AIC without posterior hippocampal novelty activation: -1981.0; AIC with posterior hippocampal novelty activation: -1985.1; BIC without posterior hippocampal novelty activation: -1964.4; BIC with posterior hippocampal novelty activation: -1963.0; model comparison: χ²[1] = 6.12, *p* = 0.013). However, the results were mixed, as the AIC values favored the model with posterior hippocampal activation, while the BIC values suggested a less clear improvement, indicating a trade-off between model fit and complexity. The model examining anterior versus posterior hippocampal activation showed that adding posterior hippocampal novelty activation did not significantly improve model fit for predicting target VTA activation compared to anterior hippocampal activation (AIC without posterior hippocampal activation: -1987.1; AIC with posterior hippocampal activation: -1986.2; BIC without posterior hippocampal activation: -1964.9; BIC with posterior hippocampal activation: -1958.5; model comparison: χ²[1] = 1.11, *p* = 0.293). The results provide some evidence of a role for posterior hippocampus novelty singling in modulating VTA engagement, although there is greater support for a specific role for the anterior hippocampus in modulating VTA responses following novelty exposure.

### Specific versus Relative Novelty

There are two ways to conceptually quantify novelty. A more specific definition may define novelty exclusively to scenes that have never been seen before, while a more relative definition may incorporate all scenes which gradually vary in their novelty signal. In this later case a familiar stimulus could still maintain a moderated novelty response which could still engage hippocampal activity. We tested whether the novelty benefit was specific to novel scenes in the preceding block (specific novelty) or a sustained signal through the block (relative novelty). To address this, we compared two models: the first model included hippocampal activation in response to novel scenes only (specific novelty), while the second model incorporated hippocampal activation in response to both novel and familiar scenes (relative novelty). Our analysis showed that the model including both novel and familiar scenes, capturing the relative novelty effect, significantly improved the prediction of subsequent VTA activation compared to the specific novelty model (AIC with specific novelty: -1609.1; AIC with relative novelty: -1616.4; BIC with specific novelty: -1587.8; BIC with relative novelty: -1589.7; model comparison: χ²[1] = 9.25, *p* = 0.002). These findings suggest that a relative novelty signal, encompassing both novel and familiar scene activations, is more critical for modulating downstream VTA engagement than a response limited to specific novel stimuli.

### Temporal Proximity to Novelty Shapes Mesolimbic Circuit Dynamics

To test whether the relationship between hippocampal novelty signals and target-related VTA activation was modulated by the temporal distance from the last novel trial, we included lag as an interaction term in the linear mixed effects model. Consistent with our hypothesis, we found a significant interaction such that the predictive effect of hippocampal novelty activation on VTA target activation decreased as lag from the last novel stimulus increased (interaction term: β = −0.0355, *SE* = 0.0157, *t*(1855) = −2.27, *p* = 0.02; Figure 7). Adding the interaction term significantly improved model fit compared to the baseline model with random intercepts only (AIC and BIC for baseline model: −1963.5, −1946.9; AIC and BIC for full model with interaction: −1970.9, −1937.7; model comparison: χ²(3) = 13.44, p = 0.004). These findings indicate that hippocampal novelty-related activation is more strongly associated with VTA target responses when the novel stimulus occurred more recently, suggesting that the modulatory influence of novelty on mesolimbic circuit engagement is temporally constrained (Figure 7).

**Figure 7.**
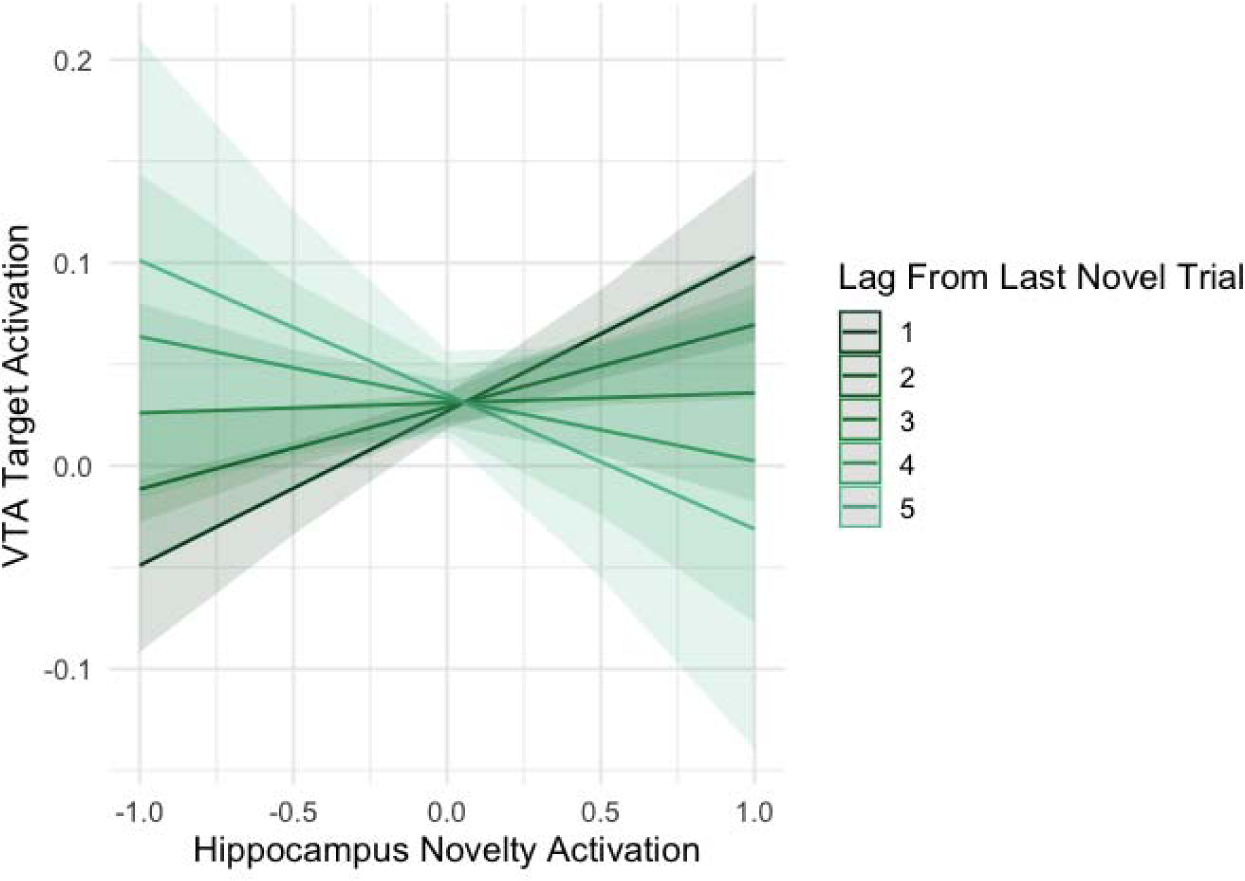
Interaction between hippocampal novelty activation and temporal lag from prior novelty in predicting VTA target activation. Estimated marginal effects from a linear mixed effects model reveal that the strength of the positive relationship between preceding hippocampal novelty activation and VTA activation during target trials decreases as the lag from the last novel stimulus increases. Lines represent estimated slopes at each lag value (1–5), with shaded bands indicating 95% confidence intervals.

### Afferent Circuit Modulation of Hippocampus Sensitivity

Rodent models hypothesize that hippocampal novelty signals upregulate VTA engagement. Critically, the VTA has a dopaminergic projection back to the VTA, increasing its sensitivity. In this way, novelty can invigorate hippocampal plasticity via VTA neuromodulatory signaling. We hypothesized that hippocampal novelty activation would predict subsequent hippocampal activation to target scenes. However, this should be mediated by the respective circuits investigated here, the hippocampal novelty circuit (hippocampus - VTA - hippocampus) and the dlPFC goal-directed circuit (dlPFC - VTA - hippocampus). We tested this using a path analysis (serial mediation model). Figure 8 depicts the effects of the paths linking hippocampal novelty activation to each mediator and hippocampal target activation, Table 1 reports the statistics. We found a significant total effect (path f, β = 0.163, 95% CI = 0.117 -0.210, *p* < 0.01), suggesting that increased hippocampal novelty activation predicts subsequent hippocampal target activation (Figure 8). Path analyses revealed that both the hippocampus-VTA novelty circuit (d*c, hippocampal novelty - VTA target - hippocampal target) and the dlPFC-VTA goal-directed circuit (b*c, dlPFC target - VTA target - hippocampal target) significantly mediated hippocampal sensitivity to subsequent targets (CFI = 1.00; RMSEA = 0.00). After controlling for these afferent circuits, the direct effect of hippocampal novelty activation on hippocampal target activation was still significant (f’, β = 0.143, 95% CI = 0.100 -0.188, *p* < 0.01).The model R² for hippocampal target activation was 0.121, indicating that afferent circuit pathways accounted for a significant portion of variance in hippocampal activation. Other indirect effects were not significant. Statistics for the other direct and indirect effects are reported in Supplemental Table 1.

**Figure 8:**
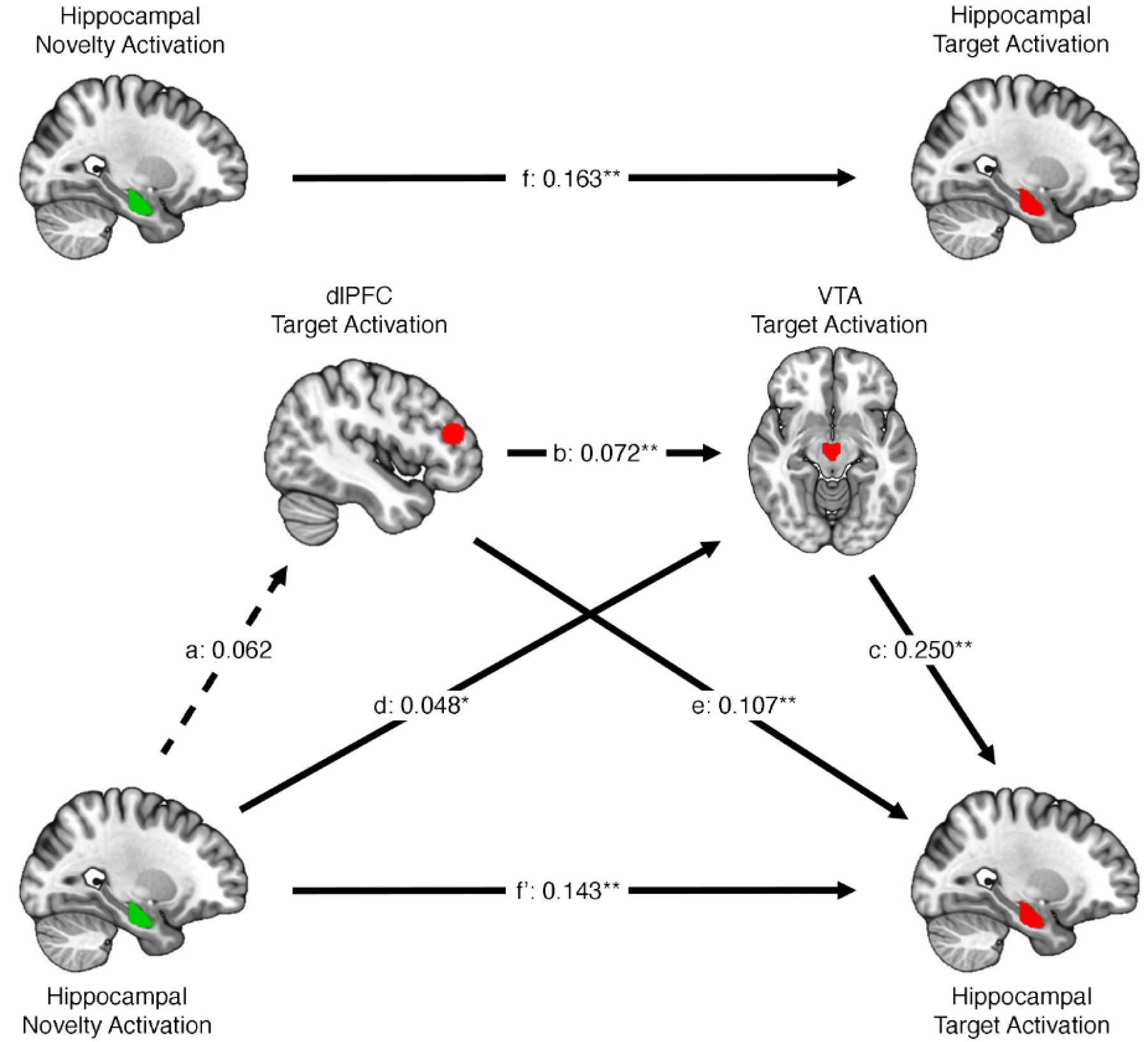
Path analysis of afferent circuits modulating hippocampal plasticity. Standardized regression coefficients are shown for paths between hippocampal novelty activation, dorsolateral prefrontal cortex (dlPFC) activation, ventral tegmental area (VTA) activation, and hippocampal activation during target trials. Solid lines represent significant paths; dashed lines indicate non-significant paths. Asterisks denote statistical significance (p < 0.05, p < 0.01).

**Table 1.** Direct and indirect effects of hippocampal (HPC) novelty activation on subsequent HPC target activation (* = *p* < 0.05, ** = *p* < 0.01)

| Outcome | $\beta$ | BootSE | BootLLCI | BootULCI |
| --- | --- | --- | --- | --- |
| **Direct Effect** |  |  |  |  |
| HPC Novelty – HPC Target | 0.143** | 0.023 | 0.100 | 0.188 |
| **HPC Novelty Circuit** |  |  |  |  |
| HPC Novelty – VTA Target – HPC Target | 0.012* | 0.005 | 0.002 | 0.021 |
| **dIPFC Goal-Directed Circuit** |  |  |  |  |
| dIPFC Target – VTA target | 0.018** | 0.004 | 0.011 | 0.026 |
| **Other Indirect Effects** |  |  |  |  |
| HPC Novelty – dIPFC Target – VTA Target – HPC Target | 0.001 | 0.001 | 0.000 | 0.003 |
| HPC Novelty – dIPFC Target – VTA Target – HPC Target | 0.007 | 0.004 | -0.001 | 0.015 |

## Discussion

Adaptive behavior requires an organism to detect and respond to changes in the environment. Novelty detection plays a crucial role in this process, acting as a signal for meaningful changes in the environment and triggering adaptive responses to goal-relevant information. However, open questions remain about how the brain translates these novelty signals to promote goal-directed target detection. Rodent research suggests that novelty can promote a positive feedback loop, wherein hippocampal novelty circuits can invigorate hippocampal responses to goal-relevant stimuli via engaging hippocampal-VTA and PFC-VTA circuits (Grace et al., 2007; Legault & Wise, 2001; Lisman et al., 2011; Lisman & Grace, 2005; Lodge & Grace, 2008). However, although rodent research provides circuit level specificity, how these circuits temporally interact during sustained motivational states (i.e. novelty) and transient goal-directed behavior (i.e., target detection) remain opaque. In the current study, we implemented a combination of advanced fMRI analysis techniques and path analysis to characterize the unique contributions of hippocampal novelty and PFC goal-directed circuits modulating VTA activation and their subsequent effect on hippocampal engagement. We found that hippocampal novelty signals and PFC goal-directed signals dynamically and synergistically predicted subsequent VTA activation. Additionally, these circuits independently mediated subsequent hippocampal engagement. These findings provide critical insights into afferent circuits modulate mesolimbic signals across time in humans and their downstream effect on hippocampal plasticity during goal-directed behavior.

### Afferent Modulation of VTA Engagement

Previous animal research has provided circuit-level framework for understanding how novelty signals affect mesolimbic engagement. Our findings dovetail prior work in animal models that describe the hippocampal-VTA loop as a crucial circuit for novelty-driven VTA engagement that unfolds across multiple temporal phases. Mechanistically, these effects should unfold in three phases: (1) Hippocampal engagement of the VTA during novelty, (2) sustained novelty-driven VTA engagement up until target image and (3) Hippocampal novelty activation predicting VTA target activation. In line with rodent models, the first phase involves direct hippocampal engagement of the VTA during novelty exposure, reflecting a disinhibitory drive of VTA activity. The present data provide clear evidence for this phase: hippocampal activation during novel scene viewing predicted concurrent VTA activation, consistent with rapid, time-locked coupling between these regions (Figure 4). This immediate coactivation establishes a foundation for subsequent novelty-to-target effects observed in later trials.

Next, we found that found that preceding anterior hippocampal novelty signals enhance subsequent VTA engagement during goal-directed target detection. Permutation analyses showed that this effect was specific to hippocampal novelty activation during the preceding trials within a block. Additionally, we observed that this effect was driven by hippocampal novelty signaling and not limbic striatum (nucleus accumbens) signaling. This result supports the notion that the anterior hippocampus plays a specialized role in detecting novelty and priming the VTA, consistent with the polysynaptic circuitry observed in non-human animals (Legault & Wise, 2001; Otmakhova et al., 2013). This homology reinforces the importance of hippocampal-VTA interactions in modulating motivational processes in dynamic environments.

A second, sustained phase of hippocampal-driven VTA modulation has been proposed to bridge novelty detection and later goal-directed responding. Although our event-related design was optimized for detecting trial-specific effects of prior novelty engagement on VTA target activity, its rapid alternation of conditions precluded the dissociation of slower, tonic VTA shifts from event-evoked responses. Nevertheless, prior work has shown that hippocampal novelty responses can elevate sustained VTA activity (Murty et al., 2017), consistent with a tonic dopaminergic mechanism that may underlie motivational readiness. Together, these findings outline a temporally nested process: rapid, phasic hippocampal engagement of the VTA during novelty that may initiate slower tonic states of elevated mesolimbic excitability, which in turn potentiate phasic VTA responses to subsequent goal-relevant events. Future work using designs with variable delays between novelty and target presentation could directly test this proposed tonic “bridge” phase, providing a more complete account of how novelty invigorates mesolimbic circuits to promote adaptive behavior.

Our findings extend prior evidence highlighting the influential role of the prefrontal cortex (PFC) in orchestrating motivated behavior through mesolimbic engagement, particularly through the modulation of phasic VTA activity. The PFC has been shown to detect goal-relevant stimuli (Blumenfeld & Ranganath, 2007; Menon & D’Esposito, 2022; Miller & Cohen, 2001), is critical for attention and executive functions (Benchenane et al., 2011; Friedman & Robbins, 2022b; Paneri & Gregoriou, 2017; Rossi et al., 2009; Zanto et al., 2011), and is implicated in motivated behavior (Cohen et al., 2014; Jimura et al., 2010; Murty & Adcock, 2014). Likewise, tasks that disrupt attention and executive function have been shown to impair motivated behavior (Elliott & Brewer, 2019). Furthermore, the PFC has been shown to stimulate the VTA in humans (Ballard et al., 2011; Murty et al., 2017), thereby potentially enhancing hippocampal sensitivity to new information (Murty & Adcock, 2014). Our finding aligns with the established role of the PFC in modulating attention and executive function to prioritize goal-relevant information. Together, these results support a model in which PFC–VTA coupling acts as a top-down mechanism linking executive functions and motivational drive to guide goal-directed behavior.

Notably, our analysis focused on a dlPFC ROI to capture the goal-oriented component of circuit dynamics, analogous to regions characterized in rodent models of motivated behavior. However, another potential contributor to the observed mesolimbic engagement may be motor readiness associated with anticipating or executing target responses. Novelty and motivational salience are known to invigorate preparatory motor processes, increasing corticospinal excitability and promoting faster behavioral responses (Bestmann et al., 2008; Lyer et al., 2010; Roesch & Olson, 2004). Additionally, the HPC-VTA circuit investigated here are part of a larger, “upward spiraling” corticostriatal and nigrostriatal circuit (Haber & Knutson, 2010). Within this framework, such motor readiness reflects coordinated interactions between limbic, associative, and motor cortico-striatal circuits: hippocampal and orbitofrontal regions project to the ventral striatum (limbic loop), which in turn communicates with dorsomedial and premotor striatal areas (associative and motor loops) via spiraling dopaminergic projections through the midbrain. These integrated loops provide a mechanism by which novelty-related hippocampal activation could invigorate mesolimbic circuits and propagate to motor planning regions to facilitate rapid goal-directed action. Future work that characterizes contemporaneous activity across premotor cortex, basal ganglia, and substantia nigra could further delineate how motivational and motor loops jointly contribute to adaptive behavior.

### Hippocampal and PFC Circuits Invigorate Hippocampal Sensitivity

Animal studies have demonstrated that dopaminergic neuromodulation is crucial for synaptic plasticity in the hippocampus (Frey et al., 1993; Frey & Morris, 1998; Huang & Kandel, 1995), increasing hippocampal sensitivity (Hammad & Wagner, 2006; Swant et al., 2008) and stabilizing hippocampal representations (McNamara et al., 2014; Martig & Mizumori, 2011; Nguyen et al., 2014; Retailleau & Morris, 2018; Tran et al., 2008; Werlen & Jones, 2015). Studies in humans have shown that increased motivational demands improve memory encoding, and that this is mediated by engagement of the VTA and hippocampus (Adcock et al., 2006; Elliott, Blais, et al., 2020; Elliott, McClure, et al., 2020; Gruber et al., 2016; Knowlton & Castel, 2022; Shigemune et al., 2014; Wittmann et al., 2007a, 2007b). Our combined use of traditional fMRI and path analysis methods allowed us to test aspects of the hippocampal-VTA-PFC network that have not been detailed in previous models, including the Lisman and Grace model. Importantly, our study provides empirical evidence for the timing and mechanistic interplay between these circuits. Specifically, we showed that the hippocampus and dlPFC independently influence VTA activity, but their effects converge to enhance hippocampal sensitivity to goal-relevant information.

These findings suggest a form of synergistic coincidence detection, where the temporal and mechanistic coordination of these signals is critical for optimal VTA modulation. Importantly, our results highlight that while the hippocampal and PFC inputs to the VTA function independently, they ultimately converge to modulate downstream hippocampal processing, as shown by our path analysis. This finding comports with rodent literature that has shown combined activation of the hippocampal novelty pathway and PFC pathways significantly amplifies both VTA neuron population activity and burst firing (Grace et al., 2007; Lodge & Grace, 2006a). This suggests that the synchronous activity of distinct afferent inputs to the VTA synergistically upregulates the VTA, which in turn invigorates the hippocampus, promoting hippocampal plasticity.

### Temporal Proximity to Novelty Invigorates VTA Engagement and Behavior

Our behavioral analyses demonstrated that temporal proximity to novelty was associated with faster goal-directed responses. Targets appearing soon after a novel stimulus were detected more rapidly, whereas increasing temporal lag from novelty predicted slower reaction times. This behavioral facilitation parallels the temporally constrained neural coupling between hippocampal novelty signals and VTA activation, suggesting that novelty exerts its greatest influence on both mesolimbic circuitry and behavior in the trials nearest to the target trial. Together, these findings indicate that novelty does not simply produce a sustained motivational state but rather creates a transient period during which hippocampal–VTA dynamics are optimized to support rapid, goal-directed responding.

### Hippocampal Long-Axis Specificity

Rodent models specify that the hippocampal novelty signal is specifically driven by the ventral hippocampus. It is also hypothesized in humans that the hippocampus is structurally and functionally heterogeneous along the anterior-posterior (long) axis. Indeed, human research has shown preferential structural (Elliott et al., 2024) and functional connectivity between the anterior hippocampus and the VTA (Cowan et al., 2021; Kahn & Shohamy, 2013). By demonstrating that anterior hippocampal novelty signals, rather than posterior, are critical for modulating VTA activity, our data add granularity to the understanding of hippocampal long-axis specificity in humans.

### Specific versus Relative Novelty

Novelty can be conceptualized as either an absolute response reserved for novel images (specific novelty) or relative, context-dependent response that scales with how unexpected a stimulus is within a given environment (relative novelty). In our task, both novel and familiar scenes were presented only once during scanning, whereas the target image was repeated many times. Thus, familiar scenes were still relatively novel within the broader task context, as they occurred infrequently and violated the dominant expectation set by the repeated target. We found that a model encompassing both preceding hippocampal activation during novel trials and hippocampal activation during familiar trials predicting target VTA activation best fit the data, supporting the influence of relative novelty. This suggests that hat hippocampal responses tracked graded contextual novelty, consistent with prior accounts proposing that hippocampal novelty responses scale continuously with stimulus expectancy and environmental statistics (Kafkas & Montaldi, 2015, 2018). Indeed, a prior study using a similar task found no hippocampal activation differences between novel and familiar scenes (Cowan et al., 2021), consistent with the possibility that familiar trials in a novel context may still carry residual novelty signals. However, these distinctions are tangential to our main question: the study was not designed to discriminate novel versus familiar stimuli or subtypes of novelty, but rather to use incidental stimuli varying in their continuous novelty to provide naturalistic variability in hippocampal–VTA circuit engagement. Future studies manipulating specific subtypes of novelty and familiarity are needed to investigate these hypotheses.

Critically, our temporal proximity results suggest that the functional impact of novelty on downstream mesolimbic engagement and behavior is most potent when novel events occur closest to target trials, producing the greatest VTA target activation and fastest goal-directed responses. This pattern is consistent with a model in which novelty not only sustains a general state of elevated hippocampal readiness but also exerts its strongest modulatory effects in the immediate aftermath of novel experiences. Rodent studies have shown that the neuromodulatory influence of novelty on synaptic plasticity not only occurs during novelty (Davis et al., 2004; Lemon & Manahan-Vaughan, 2006; Kemp & Manahan-Vaughan, 2004) but also extends beyond the initial period of exploration. Specifically, research has shown that rats exploring a novel spatial environment experience a temporally extended reduction in the threshold for long-term potentiation (LTP) induction in the hippocampal CA1 region, persisting 15 to 30 minutes after the novelty experience (Li et al., 2003; Straube et al., 2003). However, the task designs of these studies, in which an animal typically explores a novel environment freely, differ significantly from our paradigm which manipulates novelty dynamically over a short timescale. Thus, our finding of a relative novelty signal predicting VTA activation comports with rodent results, as it suggests that novelty-induced neuromodulation could continue to enhance hippocampal plasticity even when familiar stimuli are encountered in the same context. This sustained activation may reflect an adaptive mechanism whereby the hippocampus maintains an elevated state of plasticity, enhancing learning and memory processes for familiar information presented within a novel context. However, our results diverge from the animal findings by demonstrating that in a dynamic context, novelty demonstrates temporal specificity. Thus, while a relative novelty signal can invigorate the VTA to an extent, the functional impact of novelty on downstream mesolimbic engagement and target detection is most potent immediately following novel events.

### Limitations and Future Directions

Although our data establish the significance of the hippocampal and PFC circuits in VTA modulation, we acknowledge that this does not represent a complete mediation of the hippocampal target responses. There are several other afferent circuits implicated in VTA modulation, including the habenula (Christoph et al., 1986; Hennigan et al., 2015; Stamatakis & Stuber, 2012), medial prefrontal cortex (mPFC; Gao et al., 2007; Gariano & Groves, 1988; Parker et al., 2011; Sevensson & Tung, 1989; Takahashi et al., 2011), and pedunculopontine nucleus (Lodge & Grace, 2006b; Lokwan et al., 1999; Sesack & Carr, 2002; Yoo et al., 2017). Future studies should investigate these additional pathways to provide a more comprehensive understanding of how VTA activity is orchestrated by multiple neural inputs.

Recent research has also indicated a role for a locus coeruleus (LC) neuronal projection to the hippocampus in coding novelty and enhancing hippocampal plasticity (Takeuchi et al., 2016; Yamasaki & Takeuchi, 2017). It has been proposed that the LC-hippocampus pathway may specifically encode responses to distinct novelty, enhancing initial memory consolidation through heightened plasticity. This process relies on the corelease of dopamine and noradrenaline, promoting robust and vivid memory encoding for stimuli encountered in discrete, novel episodes (Duszkiewicz et al., 2019). In this way, an LC-hippocampus pathway may be better suited to encode novelty in a trial-specific manner. However, human research investigating LC-hippocampal projections and novelty coding is methodologically challenging and lacking.

Animal models of the hippocampal-VTA circuit posit that the hippocampus regulates VTA dopamine neurons. However, the VTA also contains glutamatergic and GABAergic cells (Yamaguchi et al., 2007; Nair-Roberts et al., 2008). fMRI relies on an indirect measure of neuronal activity (i.e., the BOLD signal) and does not allow for the direct classification of cell type. BOLD signals in the VTA likely reflect the integrated activity of these neuronal subtypes, complicating the interpretation of whether observed activity specifically reflects dopaminergic processes or the broader functional role of the VTA. As such, future studies combining BOLD imaging with complementary techniques (e.g., pharmacological manipulations) are needed to disentangle the contributions of different neurotransmitter and neuromodulatory systems to novelty-driven VTA-hippocampal interactions.

Although our findings demonstrate novelty-driven increases in hippocampal sensitivity, an important limitation is the absence of individual difference metrics or measures explicitly linking this neural activity to memory outcomes (probed by either self-report or explicit memory tests). While the observed hippocampal responses suggest potential for enhanced plasticity, it remains unclear whether these changes lead to measurable memory benefits. Without behavioral assessments, we cannot determine whether the novelty-driven hippocampal sensitivity observed in this study facilitates memory encoding or consolidation. To address this, future research should incorporate explicit memory testing alongside neural measures to directly evaluate the functional impact of novelty-driven VTA-hippocampal interactions. Such an approach would provide a more comprehensive understanding of the relationship between novelty, hippocampal activity, and memory, helping to bridge the gap between observed neural dynamics and their potential behavioral relevance.

Beyond the theoretical contribution to understanding hippocampal–VTA–PFC interactions, these findings also have translational relevance to understanding maladaptive behaviors and psychiatry disorders. Dysregulation of novelty processing and mesolimbic engagement has been implicated in neuropsychiatric conditions such as schizophrenia, depression, and substance use disorders, where motivational and memory processes are often impaired (Grace, 2012; Bagot et al., 2015; Keleta & Martinez, 2012). By identifying temporally specific pathways through which novelty invigorates VTA activity and downstream hippocampal sensitivity, our results provide a framework for investigating these circuits in clinical populations to better characterize disease and inform future interventions. Methodologically, the lag-based fMRI approach employed here offers a tool for probing time-dependent neuromodulatory effects in humans, which could be adapted to study circuit dysfunction in clinical populations.

## Conclusions

Our study sheds light on the complex interplay between novelty and goal-directed circuits in the brain, emphasizing the independent yet convergent roles of the hippocampus and prefrontal cortex in modulating VTA activity, promoting hippocampal plasticity to support future adaptive responses. This study highlights the dynamic coordination of distributed neural circuits in optimizing adaptive behavior. These findings have broad implications for understanding how the brain prioritizes and processes information in dynamic environments, contributing to adaptive and maladaptive behavior.

## Resource Availability

### Lead Contact

Blake Elliott

### Data and Code Availability

The data and code that support the findings of this study are openly available at https://github.com/blelliott23/

## Acknowledgments

This work was supported by the National Institute of Health (R01 MH112613 to L.E. and V.M.; R01 DA055259 & K01MH111991 to V.M.)

---

**Supplemental Table 1.** Direct and indirect effects of hippocampal (HPC) novelty activation on subsequent HPC target activation.

| **Outcome** | **$\beta$** | **BootSE** | **BootLLCI** | **BootULCI** |
| --- | --- | --- | --- | --- |
| Novel HPC - Target dlPFC (a) | 0.062 | 0.036 | -0.008 | 0.135 |
| Target dlPFC - Target VTA (b) | 0.072 | 0.014 | 0.044 | 0.100 |
| Target VTA - Target HPC (c) | 0.250 | 0.026 | 0.200 | 0.300 |
| Novel HPC - Target VTA (d) | 0.048 | 0.019 | 0.010 | 0.085 |
| Target dlPFC – Target HPC (e) | 0.107 | 0.017 | 0.071 | 0.138 |
| Novel HPC - Target HPC (f) | 0.143 | 0.023 | 0.100 | 0.188 |
| indirect_1 - a*b*c | 0.001 | 0.001 | -0.001 | 0.003 |
| indirect_2 - a*e | 0.007 | 0.004 | -0.001 | 0.015 |
| indirect_3 - d*c | 0.012 | 0.005 | 0.002 | 0.021 |
| indirect_4 - b*c | 0.018 | 0.004 | 0.011 | 0.026 |
| Total Effect - a*b*c+a*e+d*c+f | 0.163 | 0.024 | 0.117 | 0.210 |
| Total Indirect - b*c+e | 0.125 | 0.018 | 0.089 | 0.158 |

